# Assessing the application of landmark-free morphometrics to macroevolutionary analyses

**DOI:** 10.1101/2024.04.24.590959

**Authors:** James M. Mulqueeney, Thomas H. G. Ezard, Anjali Goswami

**Affiliations:** Department of Ocean & Earth Science, National Oceanography Centre Southampton, University of Southampton Waterfront Campus, Southampton, UK; Department of Life Sciences, Natural History Museum, London, UK

## Abstract

The study of phenotypic evolution has been transformed by methods allowing three-dimensional quantification of anatomical form. The present state-of-the-art 3D geometric morphometrics, which relies heavily on manual landmarking, is time-consuming, prone to operator bias, and cannot effectively compare disparate shapes. Emerging automated approaches, notably landmark-free techniques, offer promise but have only been applied to closely related species. Here, we compare landmark-free approaches with high-density geometric morphometric methods on 322 mammals across 180 families. Leveraging the benefits of Poisson meshes, which combine open and closed projections, we show how landmark-free methods have greater power to differentiate major taxonomic groups. Although there is coarse correspondence in shape variation between the two methods, the finer features of the landmark-free approach likely reflect its broader sampling of the surface structure. Our study underscores the robustness of landmark-free methods for large-scale comparative analysis and helps propel morphometrics into a new era of bigger data.

---

Morphometrics, the quantitative analysis of shape, is a well-established family of methods in the field of biology^1^. In recent decades, geometric morphometrics has emerged as the gold standard for addressing evolutionary questions of shape in diverse datasets^2^. Typically, this approach relies on the manual placement of landmarks to produce two or three-dimensional coordinates by labelling homologous anatomical loci^3,4^. Raw coordinates are then transformed using methods such as Procrustes superimposition^5^ to register objects to a common frame and thereby isolate biological variation.

Despite advancements to incorporate high-density morphometric analysis^6^, including semi-automated placement of sliding semilandmarks landmarks^7^, geometric morphometric methods remain largely manual and thus are time-consuming, prone to observer bias, and lack repeatability^8^. With the increasing accessibility and affordability of high-resolution imaging^9,10^, coupled with the development of tools for automated image segmentation^11,12^, databases of 3D images are expanding, providing vast amounts of data for morphometric analysis^13,14^. Thus, there is now a pressing need to improve the software and methodology we use to analyse morphological evolution.

In addition to speed and repeatability, the requirement of homology for landmark placement, while important for meaningful comparability across specimens, limits not only processing time but their applicability when comparing disparate taxa, as identifiable homologous points become more obscure and fewer in number, even within homologous structures^15^. Consequently, the reduction in the number of discernible landmarks when analysing phylogenetically distinct taxa results in the capture and comparison of only a minimal amount of variation, leading to weaker biological inferences^16^.

New automated approaches, such as “landmark- or homology-free” morphometric methods have been developed to overcome these issues. One example is diffeomorphic methods, which utilise deformable grids and registered surface points known as control points to map samples onto a standardised atlas of mean shape^17^. While these methods have shown success at analysing variation within species^18^, their effectiveness at comparing shape at higher taxonomic levels and their ability to capture comparable aspects of variation without being tied to homologous points remains untested.

In this study, we compare the estimation of shape variation using landmark-free methods with those using manual landmarking and semi-landmarking techniques (hereafter referred to as manual landmarking for brevity). Specifically, we employ a form of the diffeomorphic methods known as a Deterministic Atlas Analysis (DAA) in the software Deformetrica^19^ by following the landmark-free pipeline developed by Toussaint et al.,^18^. We apply these methodologies to 322 crania of crown and stem placental mammals to evaluate the speed, repeatability, and accuracy of each approach.

Using a two-way partial least squares analysis, we directly compare the results over a range of number of control points by adjusting the kernel width parameter, which determines the spatial extent of neighbouring data points considered in the analysis. This adjustment aims to identify the optimal number of control points using a DAA. Additionally, we assess the impact of mesh topology on the results by including both computed tomography (CT) and surface-generated meshes in our analyses. Our findings demonstrate that, once modality issues are overcome, the DAA method significantly expands spatial coverage of morphology and substantially reduces processing times compared to manual landmarking techniques, thereby paving the way for a transformative shift towards ’Big Data’ opportunities within the field of morphometrics.

## Results

### Comparison of aligned-only and Poisson meshes

Adjusting the kernel width alters the spatial extent of neighbouring data points considered in each analysis. Here we used kernel widths of 40.0mm, 20.0mm and 10.0mm in the DAA, yielding 45, 270 and 1782 control points, respectively (Fig. 1). Noteworthy disparities emerged between the results obtained from the aligned-only (mixed modalities) and the standardised filled Poisson meshes aligned using landmarks (Fig. 2, see methods). A DAA of the aligned-only data identified an artificial distinction between specimens with closed and open surfaces, i.e., surface scans and micro-CT scans, respectively (Fig. 3, Extended Data Fig. 1 and Supplementary Tables 1-3). Specimens with unfilled surfaces predominantly clustered toward the positive end of principal component (PC) 1, while those with closed surfaces were located towards the negative end. Standardising the data by closing the meshes and applying a Poisson distribution mitigated the influence of mesh type, with the results showing considerable phylogenetic structure compared to the non-standardised meshes (Fig. 4, Extended Data Fig. 2 and Supplementary Tables 4-6).

**Fig 1.**
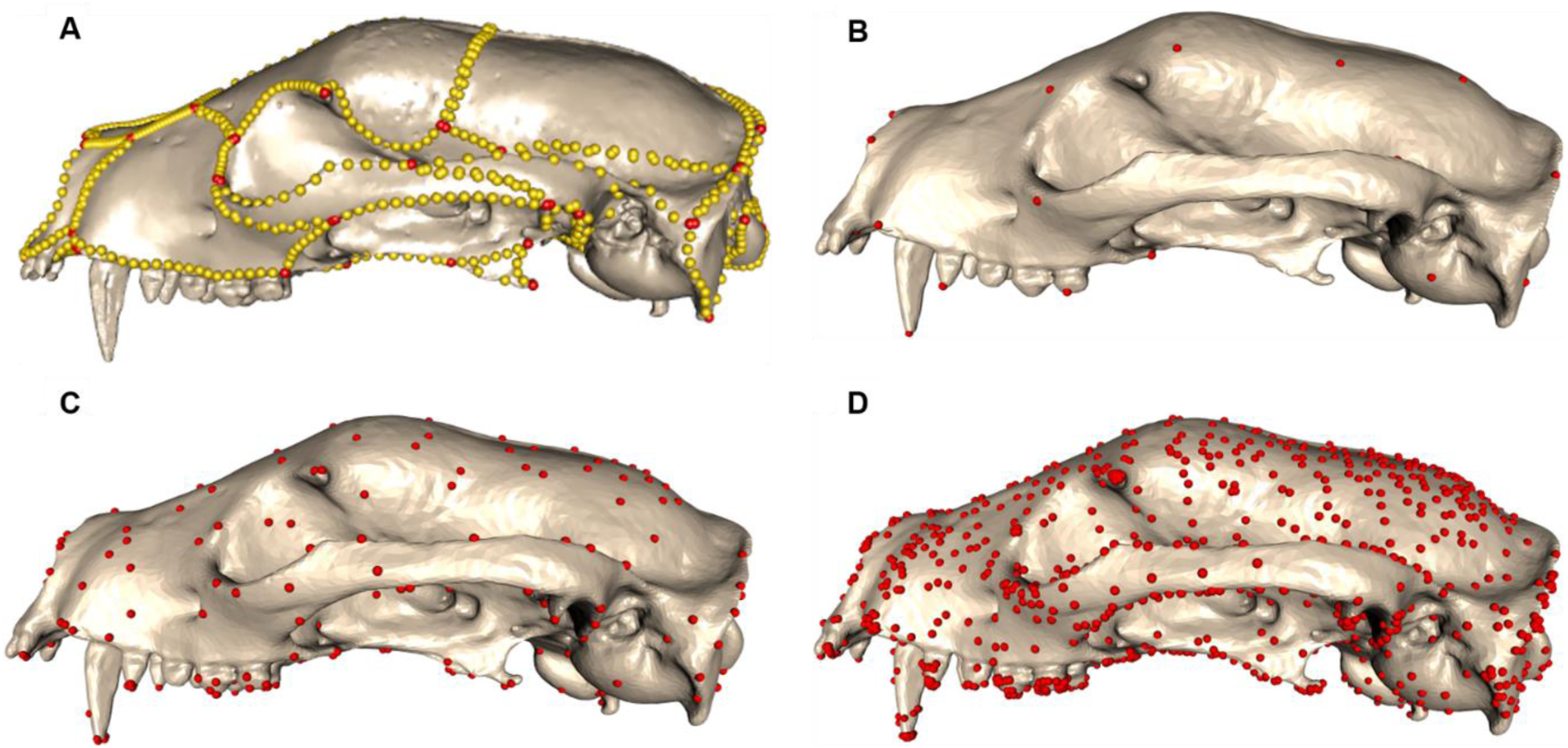
Comparison of manual landmarking scheme with a range of control points. An assessment of morphometric methods is conducted on the 3D mesh of the mammal skull, specifically here on *Arctictis binturong* (MNHN 1936-1529). The comparative analysis encompasses (**a**) a manual landmarking approach involving 754 landmarks and sliding semilandmarks, contrasted with control points derived from a Deterministic Atlas Analysis (DAA) utilising (**b**) a kernel width of 40.0mm with 45 control points, (**c**) a kernel width of 20.0mm with 270 control points, and (**d**) a kernel width of 10.0mm with 1782 control points.

**Fig 2.**
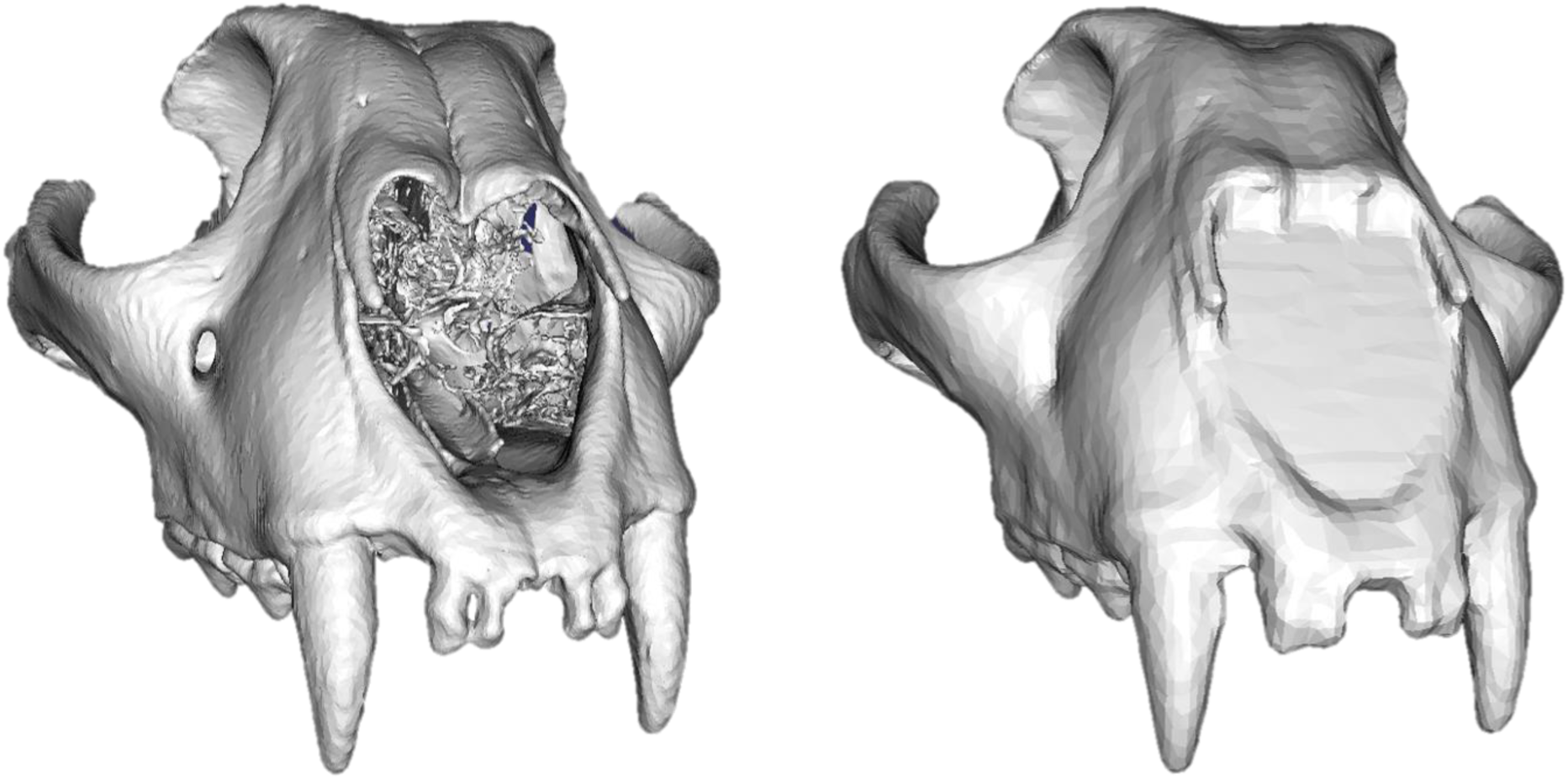
Generating Poisson meshes. Processing the mesh involves filling in holes and redistributing faces and vertices through the application of a Poisson distribution. This is demonstrated in the mesh generated from CT data, as illustrated in the case of *Acinonyx jubatus*.

**Fig 3.**
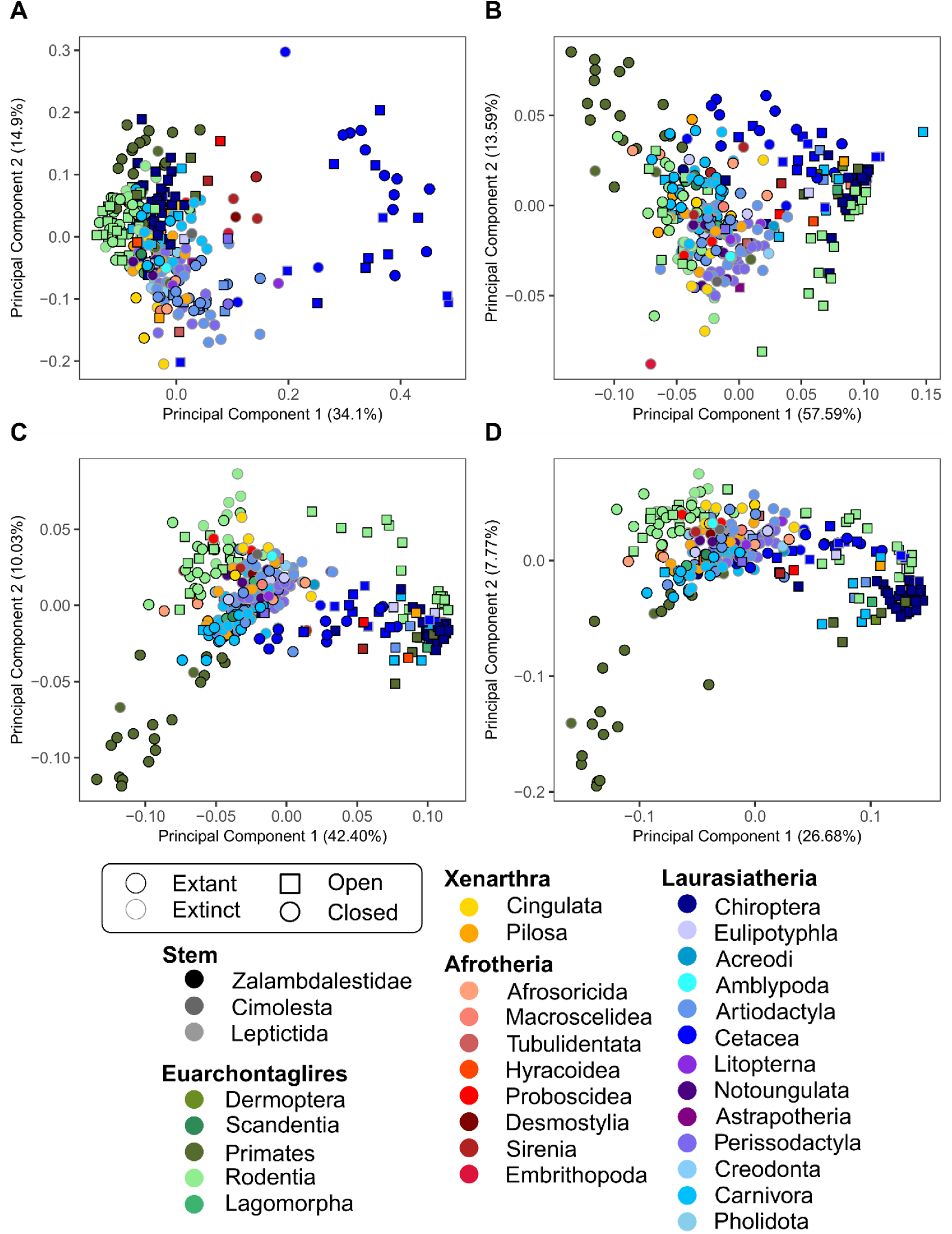
Comparison of manual landmarking and Deterministic Atlas Analysis (DAA) using aligned-only data. Cranial morphospace for placental mammals with aligned-only specimens along axes PC1 and PC2, comparing the results between (**a**) manual landmarking with 754 landmarks and sliding semilandmarks and DAA method using (**b**) a kernel width of 40.0mm resulting in 45 control points, (**c**) a kernel width of 20.0mm resulting in 270 control points and (**d**) a kernel width of 10.0mm resulting in 1782 control points.

**Fig 4.**
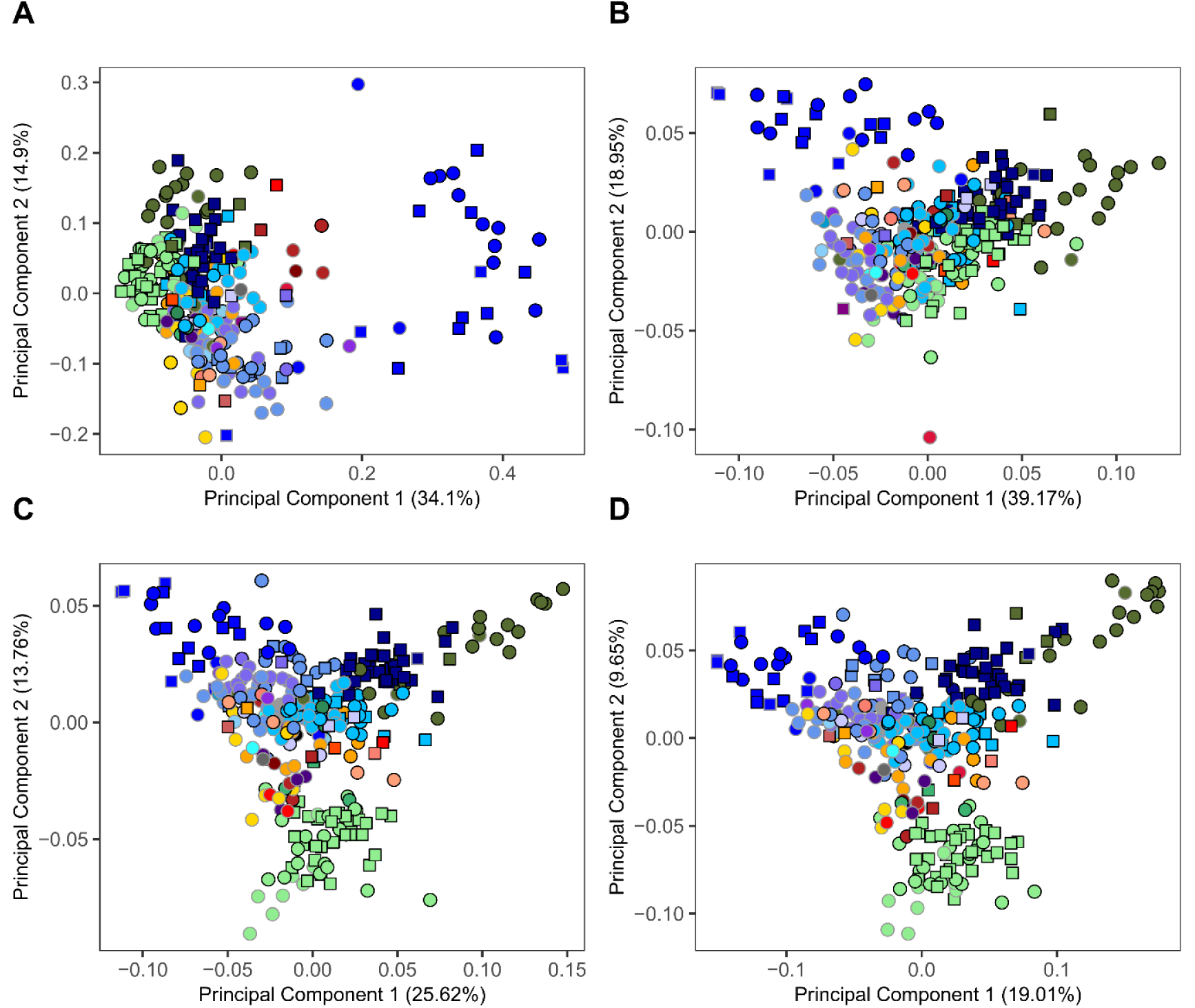
Comparison of manual landmarking and Deterministic Atlas Analysis (DAA) using Poisson meshes. Cranial morphospace for placental mammals with Poisson distributed specimens along axes PC1 and PC2, comparing the results between (**a**) manual landmarking with 754 landmarks and sliding semilandmarks and deterministic atlas analysis using (**b**) a kernel width of 40.0mm resulting in 45 control points, (**c**) a kernel width of 20.0mm resulting in 270 control points and (**d**) a kernel width of 10.0mm resulting in 1782 control points.

The partial least squares (PLS) analysis revealed a significant shift in distribution between aligned-only and filled Poisson meshes (Fig. 5). While all PLS results exhibited a significant correlation between the results of the manual landmarking and the DAA (p<0.001; 1000 permutations; Extended Data Table 1), a marked enhancement in the mean correlation was observed with the utilisation of Poisson meshes (M=0.884, SD= 0.00462), in contrast to the aligned-only data (M=0.558, SD= 0.0655), t (2.02) = 8.618, p<0.05).

**Fig 5.**
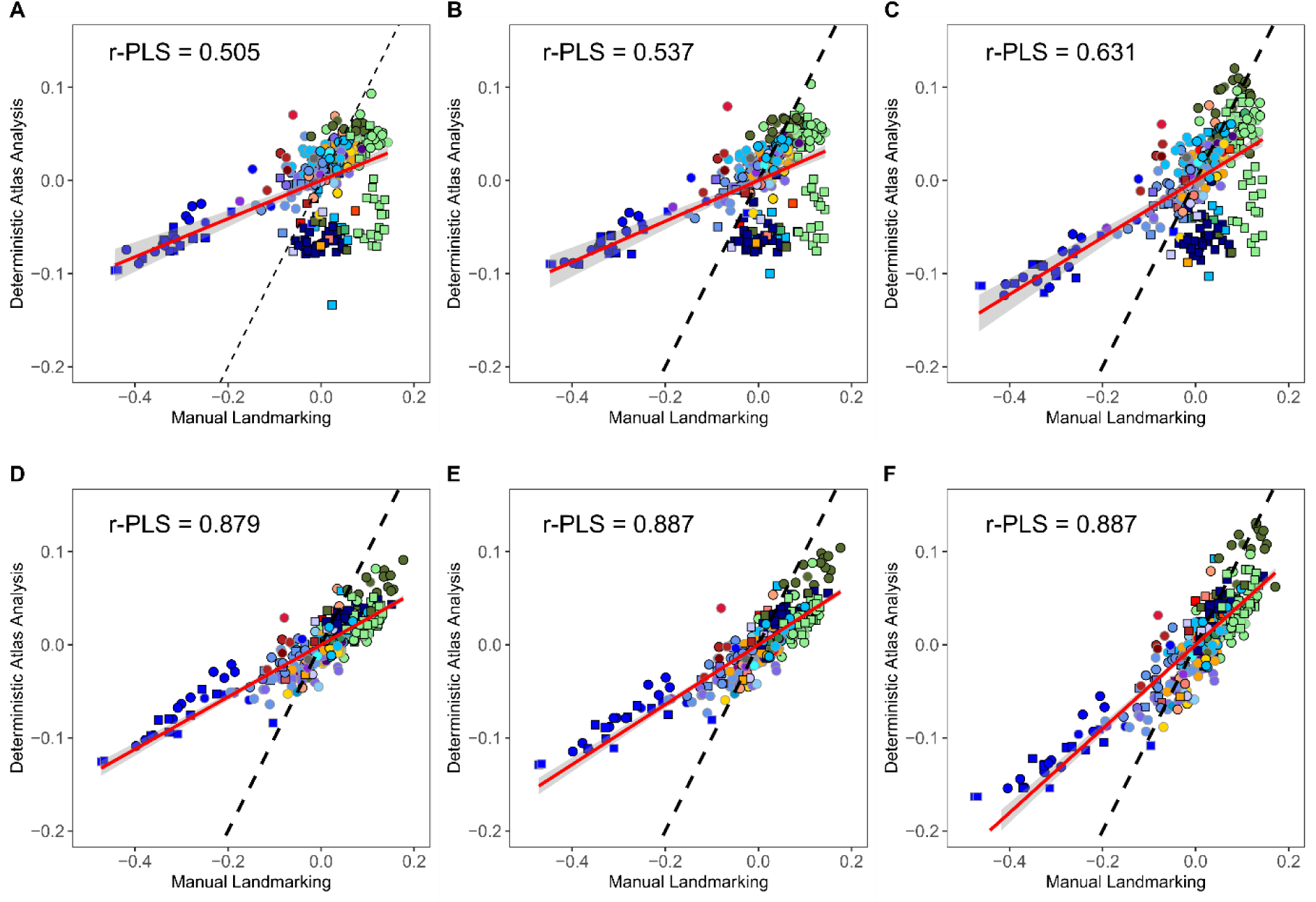
Regression plots for the partial least squares (PLS) comparison between manual landmarking and Deterministic Atlas Analysis (DAA). Comparison of the aligned-only data for (a) a kernel width of 40mm with 45 control points, (b) a kernel width of 20mm with 270 control points, (c) a kernel width of 10mm with 1782 control points and for the Poisson meshes for (d) a kernel width of 40mm with 45 control points, (e) a kernel width of 20mm with 270 control points and (f) a kernel width of 10mm with 1782 control points. The red line shows the correlation between the values and the dashed line indicates a line of equality (x=y). All r-PLS results are significant (p <0.001).

In the aligned-only data, points deviating furthest from the line of best fit tended to realign closer to both the regression line and the line of equality (x=y) when analysed within the context of Poisson meshes (Fig. 5). This realignment predominantly impacted the open meshes, which results in the improved correlation between the manual landmarking and DAA (see Fig. 5; Extended Data Table 1). Particularly noteworthy is the discernible impact on orders such as bats and rodents, characterised by a higher proportion of CT meshes. This finding aligns with the trends elucidated in the PC plots, thereby further underscoring the efficacy of Poisson meshes in refining analytical outcomes for these taxa. Consequently, all subsequent comparisons were exclusively conducted using the Poisson meshes.

### Comparison between manual landmarking and landmark-free methods

The results obtained from employing Poisson meshes in the DAA (Supplementary Table 4-6) and manual landmarking analysis (Supplementary Table 7) demonstrated a robust correlation across all three sets of control (p<0.01, 1000 permutations), with minimal or negligible enhancement as the number of control points increased (Extended Data Table 1). When controlling the number of PC axes to 321 for both the manual landmarking and the DAA we found significant correlations between the percentages of variation controlled by each axes across all control points (see Extended Data Table 2, Supplementary Tables 8-13).

In exploring the morphospace patterns, we unveil distinct similarities and disparities between the manual landmarking and DAA PCA plots (see Fig. 4, Extended Data Fig. 2). Rodents displayed distinct clustering across the DAA, but, with 45 control points, this distinction manifested primarily along PC3, akin to the outcomes of manual landmarking. Whereas with 270 and 1782 control points, by contrast, rodents occupied the most negative values along PC2. Similarly, other clades such as carnivorans and artiodactyls were predominantly concentrated within a single region of the DAA morphospace, consistent with the outcomes of manual landmarking, albeit being more distinct along PC1 as opposed to PC2. Cetaceans consistently exhibited a notable separation from other orders, occupying extreme positive values in manual landmarking and negative values in DAA for PC1. However, this separation was more pronounced in the manual landmarking approach.

Across the initial four PC axes, the taxonomic orders were more distinctly discernible when employing DAA (except for cetaceans), with a more pronounced clustering of species within their clades. This distinction was particularly evident when employing a larger number of control points, which facilitated improved spatial coverage across the cranium. These trends were further elucidated in the PLS plots, wherein cetaceans notably deviated the correlation from the line of equality (x=y) but began to shift closer when using more control points (see Fig. 5). Conversely, all other orders tended to exhibit a more precise correlation across all control points with some minor improvements as the number of control points increased.

### Comparisons of estimated mean shape using heatmaps

Visualisation of shape variation on the left side of the cranium using heatmaps for the specimen of *Cacajao calvus* revealed strong similarities in the magnitude and location of captured variation between the manual and automated approaches (Fig. 6). Negative displacement was most pronounced in the parietal region of the cranium across all analyses, though marginally smaller in the manual landmarking method compared to the DAA method. Conversely, positive displacement was prominently observed in the eye sockets across both methods, albeit more distinctly in the DAA method.

**Fig 6.**
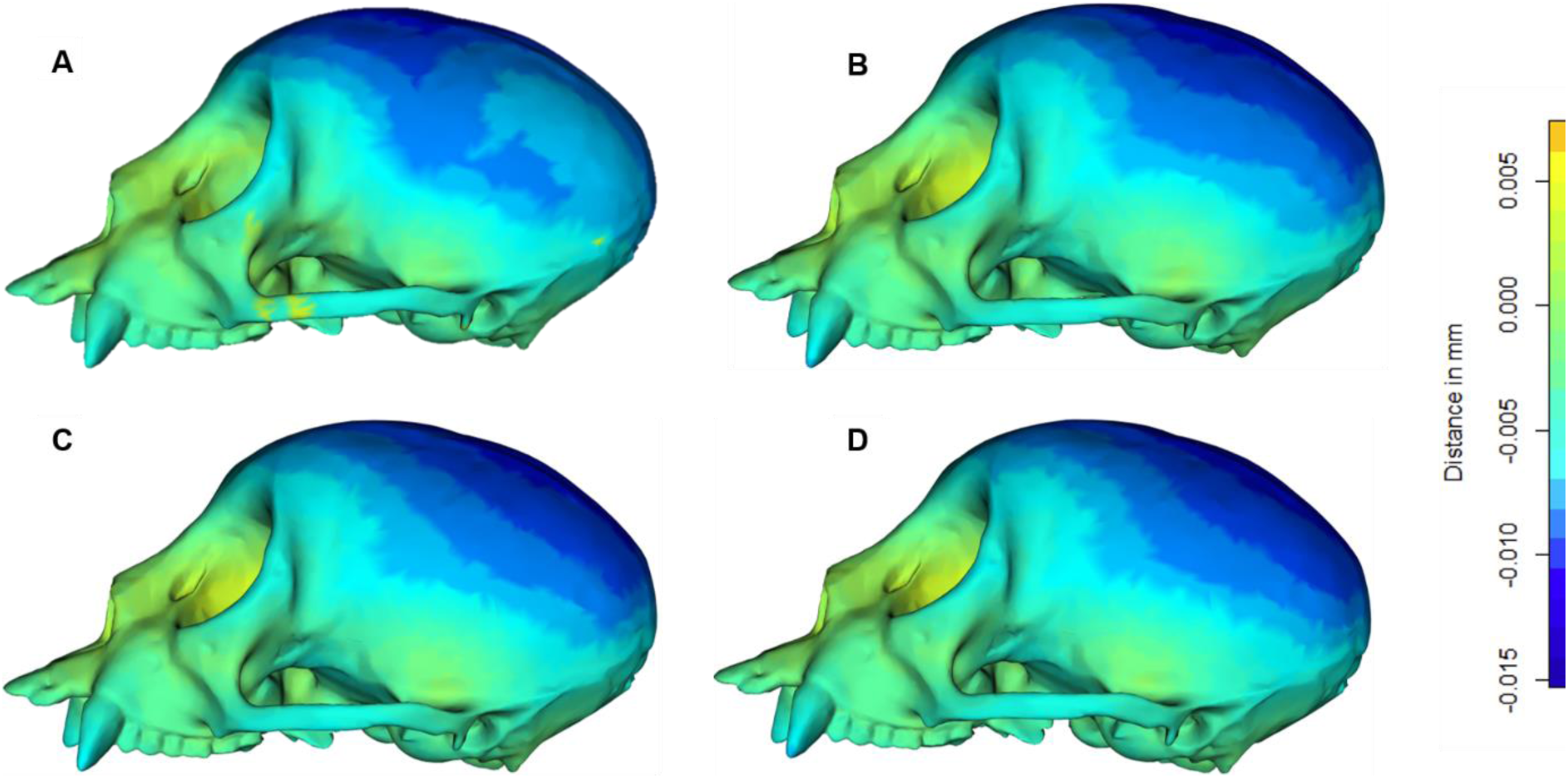
Comparing estimates of mean shape using heatmaps. Displacement heatmaps produced by superimposing the mean shape of the entire 322 specimens against the atlas specimen, *Cacajao calvus* (NHMUK ZD 1928.4.27.6) for (**a**) manual landmarking scheme with 754 landmarks and sliding semilandmarks compared to control points generated using a Deterministic Atlas Analysis (DAA) with, (**b**) a kernel width of 40mm with 45 control points, (**c**) a kernel width of 20mm with 270 control points and (**d**) a kernel width of 40mm with 1782 control points.

## Discussion

To propel the field of morphometrics into a new era of big data, it is imperative to pioneer methodologies that rapidly acquire precise high-dimensional morphological data across vast and varied datasets. This study underscores the transformative potential of the landmark-free Deterministic Atlas Analysis (DAA) in achieving this goal. The approach seamlessly scales from intraspecific datasets^18,19^ to large, diverse datasets to facilitate phylogenetic comparisons and enhance downstream evolutionary analyses, thereby offering a more accurate depiction of the complex evolutionary processes underlying the observed phenotypic variation^20^.

Not only does the DAA capture morphological variation with a robust correlation to manual landmarking but also surpasses it in terms of accuracy, efficiency, and scalability. This superiority is manifested through increased spatial coverage with the DAA approach circumventing the constraints associated with comparing morphology solely based on homologous points. As a result, we achieve greater discriminative power between taxonomic clades using this landmark-free method, with perhaps the exception of cetaceans. Moreover, the efficiency gains are substantial; processing all 322 meshes with the DAA method took ∼120 hours, significantly less than the ∼750 hours of expert time required for manual landmarking.

Our study also provides a substantial methodological advancement in the application of the DAA in the form of standardised Poisson meshes. This innovation facilitates the use of mixed modalities (micro-CT and surface scans) in the same analyses whilst maintaining a high correlation with manual landmarking results. Together, these findings underscore the pivotal role of landmark-free approaches like DAA in advancing morphometrics and ushering in a new era of data-driven research in the field.

The results suggest that, unlike manual landmarking methods, the DAA approach used in this study is influenced by variability between the modalities of the mesh rather than solely by shape. Prior research has underscored the importance of using surfaces with uniform geometry and topology for obtaining accurate results when using landmark-free methods^21,22^. In our case, the mixture of surface and CT scans, used in the analysis, led to the inclusion of meshes with both closed and open surfaces, which strongly impacted the results. As a result, analysing datasets with a mix of micro-CT and surface scans poses challenges, with the findings highlighting the need for standardised data, either using fully open or closed specimens (rather than mixed) depending upon the data available. In our case, we created filled Poisson meshes from open surface meshes to standardise surfaces for analysis.

Though the manual nature of this process made it time-consuming and less repeatable, it remains significantly quicker than manual landmarking. In the future, the mesh filling process is likely to be optimised further, benefiting from the application of machine learning approaches^23^ and increased volumes of high-quality training data^12^, which will alleviate the current bottleneck.

The findings of this study reveal a significant correlation between the results acquired through landmarking techniques and the DAA, suggesting that when appropriately applied, landmark-free methods can effectively measure shape akin to manual landmarking methods. However, notable differences emerged between the two methods.

When using the landmark-free method, we more precisely captured the classic dolichocephalic to brachycephalic trend, which was prevalent along PC axes 1, supporting results obtained using linear morphometrics^24^. In contrast, this aspect of shape variation dominates PC2 using manual landmarks, but we see a less pronounced separation between clades, particularly when compared to the DAA when using a larger number of control points. This disparity is likely due to the increased spatial sampling across the entire cranium facilitated by the DAA, which enables more accurate quantifications of shape variation^7^.

The manual landmarking scheme used here includes landmarks and sliding semilandmarks placed on sutures between cranial elements (Fig. 1a). This allows measurements of the relative positions and proportions of elements within the cranium, something not currently possible using the landmark-free approach. However, this also leads to far patchier coverage of the skull (Fig. 1) and to an unwanted bias in pre-selecting where variation is likely to occur^25^. In the heatmaps (Fig. 6) we observe greater displacement across the cranium using the landmark-free results, which may suggest these changes in distribution affect the results. Notably, the regions highlighted in the heatmaps are precisely those that suffer from a paucity of homologous landmarks, such as the cranial vault, and, as a result, are generally poorly represented in geometric morphometric analyses. Conversely, the absence of element segregation in the DAA may lead to decreased differentiation of cetaceans due to the failure to capture their distinctive features, such as significant alterations in the relative positions of cranial elements and the absence of midline convergence of parietal bones within the cranial vault.

Though some manual landmarking studies can densely sample across the cranium using landmarks and patches^26^, and thus overcome this issue of data paucity, they remain difficult to reproduce and are extremely time-consuming and hence are rarely applied to such large datasets. Moreover, using these techniques is also less useful when comparing disparate taxa, as there are fewer landmarks available to capture variation^7^. Therefore, the DAA method is beneficial in that it helps liberate the need for homology when comparing shape, a major limiting factor in how morphological variation is often compared ^27^, provided a sensible number of control points to sufficiently capture variation and biological signal without overfitting is used^28^.

Alongside the better representation of shape, a primary benefit of the landmark-free approach is a substantial improvement in processing times. Ignoring the need to standardise the mesh data, the DAA method took under 8 hours to run (with minimal differences across kernel widths), compared to ∼750 hours for full-time manual landmarking by an expert on mammal cranial morphology, not including additional time for error checking. These differences would become even more pronounced as the number of specimens and landmarks increases, meaning the efficiency gains improve as data resolution increases.

Currently, landmark-free methods may face challenges that have been resolved for manual landmarking techniques, including scenarios with incomplete specimens or absent structures. Manual landmarking analyses have developed strategies to mitigate these issues, such as employing missing data estimations^29^ or interpolating missing landmarks^27^. The absence of these techniques in landmark-free methods will impact results, particularly when assessing fine-scale morphological changes^30^. Notably, within our analysis, specimens with missing data can occupy extreme regions of morphospace, as observed in *Euhapsis ellicottae* (Fig. 4d) when using a larger number of control points. Consequently, there is a pressing need to address the influence of missing features in landmark-free techniques. While not explored in this study, novel approaches using deep learning could potentially impute missing regions of morphology^31^.

The landmark-free method also included anatomical features typically excluded in manual landmarking analyses. Within this dataset, these features include horns and teeth. Particularly when using a lower number of control points, the presence of such features impacts the placement of specimens within morphospace. For instance, the horns of *Arsnotherium zitelli* and the teeth of *Odobenus rosmarus* lead to these specimens occupying distinct areas of morphospace (Fig. 4b). Although the influence of these structures decreases with a higher number of control points, devising methods to mitigate these effects remains advantageous. Emerging techniques, such as MeshCNN which allows for mesh parcellation, hold promise in accurately and efficiently removing these objects from meshes^32^.

Unlike landmarks, the current implementation of the DAA employed in this study cannot parcellate the mesh into individual elements. Consequently, investigations into how distinct elements evolve independently^33^ and their interrelationship through phenotypic integration and modularity^34,35^ are hindered. Fully harnessing the potential of these methods requires developing accurate and efficient ways to parcel and separate anatomical regions. Borrowing from landmarking techniques, control points could be allocated to specific regions, or distinct anatomical regions could be identified and individual meshes generated. Techniques in automated image segmentation^36,37^ could be adapted to be used on the voxelised meshes or CT image data. Alternatively, the mesh could be parcellated using methodologies akin to those proposed for removing anatomical regions^32^, potentially offering a more suitable approach. Such advancements are essential to fully realise the benefits of using landmark-free methods.

Transitioning to landmark-free methods would represent a transformative shift in the field. By helping eliminate both inter- and intra-operator errors associated with manual landmark placement, these methods significantly enhance repeatability and mitigate bias in shape variation comparisons^38,39^. Notably, the DAA method, as applied in this study, offers a straightforward implementation process and delivers remarkable efficiency gains, substantially reducing processing times compared to manual landmarking methods. Therefore, this method may be better suited than other landmark-free methods such as GPSA^40^ and auto3dgm^41^, which do not offer the same accuracy and efficiency in the alignment of meshes. In an era marked by the proliferation of phenotypic data, the ability to swiftly and accurately analyse morphological variation becomes pivotal in leveraging the gains of modern imaging tools^20^. Therefore, the time-saving benefits inherent to approaches like landmark-free methods hold immense value when handling larger phenotypic datasets, enabling more expansive and robust comparisons of shape.

The most significant and pivotal advancement afforded by the DAA method resides in its unparalleled capacity to enable high-resolution shape comparisons, especially in scenarios where specimens lack an abundance of identifiable homologous points. Currently, capturing shape using manual landmarks means comparisons are rooted in the need for homology, thereby limiting effective comparisons of intricate and diverse shapes^25^. Applying landmark-free methods circumvents this problem, enabling seamless comparisons of shape throughout the entire structure, even when homology is limited or absent^42^. As a result, the application of DAA liberates comparisons across disparate taxa from the constraints of being confined solely to small homologous points, instead allowing for comprehensive comparisons across the entirety of the structure.

These improvements offered by landmark-free methods in analysing extensive and diverse datasets represent a monumental leap forward for the study of evolutionary and comparative morphology. They allow shape analyses to keep pace and match the improvements in 3D image data availability in recent years. As a result, comparative and evolutionary studies can be vastly scaled up, heralding the advent of ’Big Data’ opportunities and marking the transition into the phenomics era within the field of morphometrics^20^.

## Online Methods

### Data collection and manual landmarking scheme

We obtained meshes and geometric morphometric data for manually collected landmarks and semilandmarks data from Goswami et al.,^43^. This dataset comprised 322 crown and stem placental mammals, including 207 extant and 115 extinct species (Supplementary Table 14).

The manual landmarking scheme consisted of 66 3D landmarks and 69 semilandmark curves collected for the left side of the skull, which was done using Stratovan Checkpoint (Stratovan, Davis, CA, USA). Landmarks and semilandmarks were imported into R v.4.3.1 for analysis, where curves were resampled to a common number of semilandmarks and slid to minimise bending energy, which measures and optimises local shape differences versus the mean shape. Generalised Procrustes analysis was then used to register the landmarks, resulting in a total of 754 3D landmarks and sliding semilandmarks (Supplementary Table 15). Principal component analysis was performed using Procrustes-aligned 3D data in R for visualisation.

### Mesh processing

To assess the impact of mesh type on the results of the Deterministic Atlas Analysis (DAA), two sets of meshes were utilised. First, we standardised the mesh data to the Procrustes-transformed data used in the manual landmarking analysis by applying Procrustes transformations to all 322 meshes using the Morpho^44^ v.2.12 package in R (Supplementary Table 16). The first set of meshes contained the Procrustes-transformed mesh data in their original state, while the second set comprised Poisson distributed meshes only. The latter were created by filling holes in the open surface meshes through voxelisation and segmentation processes in Dragonfly (Object Research Systems, Canada), followed by the redistribution of faces and vertices using a Screen Poisson distribution^45^ in Meshlab^46^ v.2023.12. Meshes were smoothed and decimated to 50,000 faces for analysis to reduce computational time while maintaining overall topology.

### Deterministic Atlas Analysis (DAA)

Data were inputted into the DAA within Deformetrica^19^ following the method outlined by Toussaint et al.^18^. The mesh of *Arctictis binturong* was used as the atlas for all analyses. Varying kernel widths of 40.0mm, 20.0mm, and 10.0mm were utilised due to their varying effects on control point generation. Keops kernel (PyKeops; https://pypi.org/project/pykeops/) was employed for its performance with large datasets. All analyses were run for 150 iterations with an initial step size of 0.01 and a noise parameter of 1.0. Non-linear kernel principal component analysis^47^ (kPCA) was used for visualisation into 321 axes to mirror the manual landmarking results, using 1000 iterations and a Gamma value of 2.5×10^−6^. All analyses were conducted using Ubuntu (22.04.2) for Windows on a workstation with an Intel(R) Core (TM) i7-7700 CPU at 3.60GHz with 64.0 GB and an NVIDIA GeForce RTX 2090 graphics card with 8.0 GB of dedicated GPU memory.

### Comparing the manual landmarking and landmark-free methods

We used the R package, geomorph^48^ v.4.05 to perform a two-block partial least squares analysis^49^ (PLS) to measure the similarity between the manual landmarking and landmark-free results. This included independent comparisons of the effect of mesh processing and the number of control points. Each PLS comparison was run for 1000 iterations with a threshold p-value of 0.001.

Shape variation was then visualised using the R package Morpho^44^. A thin-plate spline was employed to morph the specimen, *Cacajao calvus,* to the mean landmark shape with the mesh displacement measured and used to create heatmaps showing areas of variation. The same method was then used to visualise the shape variation in the DAA approach by comparing the *C.calvus* to the estimated mean shape atlas generated from each of the independent analyses.

### Data accessibility

All final raw data used in the paper are available in the Supplementary Material and at https://github.com/JamesMulqueeney/Deterministic-Atlas-Analysis. All code used in the paper is also stored in this GitHub repository. 3D meshes for all specimens are available for free download on phenome10k.org or morphosource.org (see data S1) unless restricted. Processed data and other large files are available in the Dryad Repository: DOI: 10.5061/dryad.xksn02vps.

### Authors’ contributions

J.M.M, T.H.G.E and A.G designed the concept of the study. A.G performed the data collection. J.M.M performed all of the analysis and statistical comparisons. J.M.M wrote the initial draft manuscript, which was iterated with A.G. and T.H.G.E. All authors provided ideas and discussion to the paper and have read and approved the final version.

### Funding

J.M.M was supported by the Natural Environmental Research Council [grant number NE/S007210/1]. J.M.M and T.H.G.E were also supported through the Natural Environmental Research Council [grant number NE/P019269/1]. A.G was supported by Leverhulme Trust grant RPG-2021-424 and European Research Council grant STG-2014-637171.

## Acknowledgements

The authors would like to thank all the other members of the Goswami and Ezard Labs who generously provided ideas for the concept of the paper.

## Extended Data Figures

**Extended Data Fig 1.**
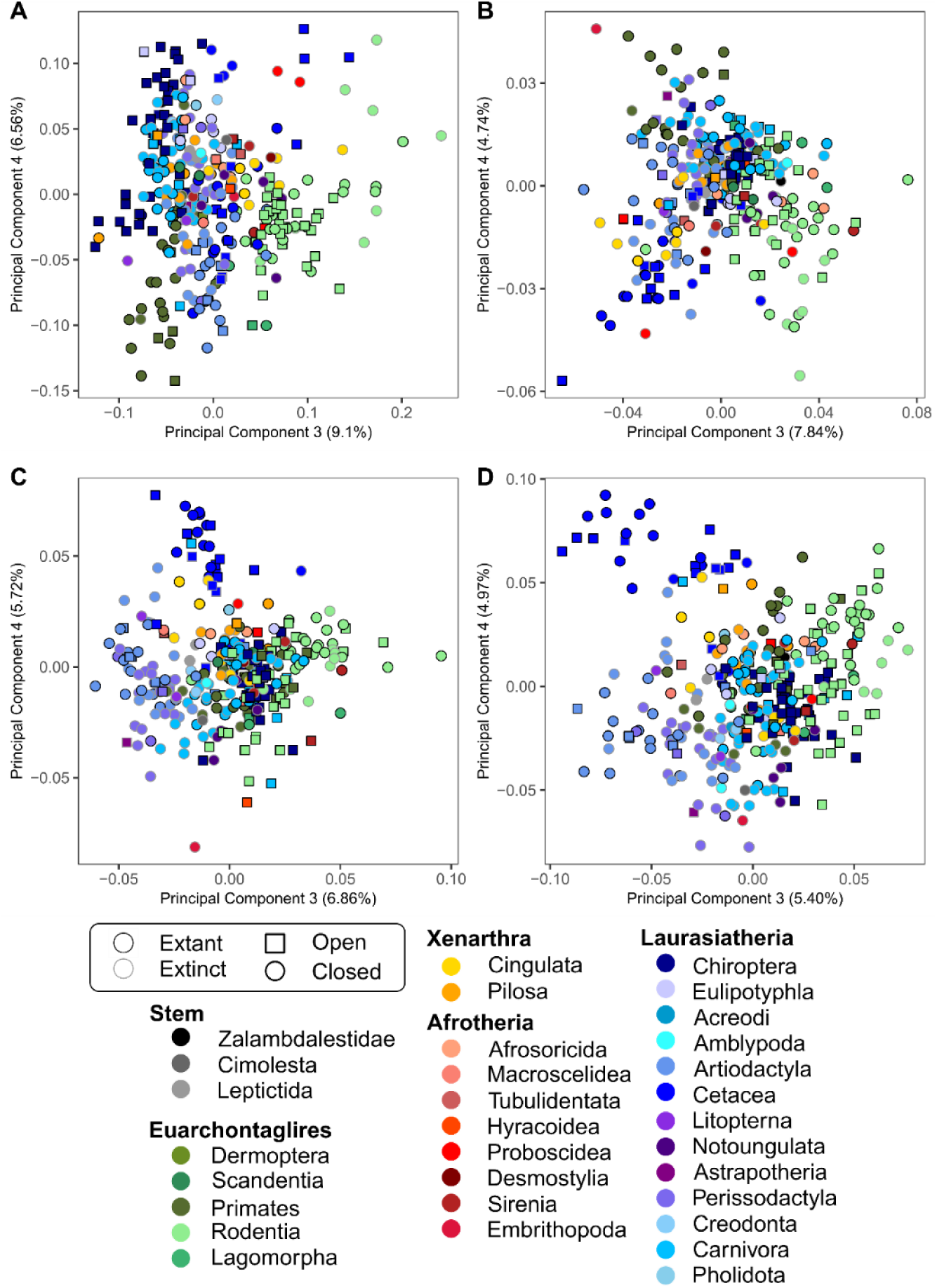
Comparison of manual landmarking and Deterministic Atlas Analysis (DAA) using aligned-only data. Cranial morphospace for placental mammals with aligned-only specimens along axes PC3 and PC4, comparing the results between (a) manual landmarking with 754 landmarks and sliding semilandmarks and deterministic atlas analysis using (b) a kernel width of 40.0 resulting in 45 control points, (c) a kernel width of 20.0 resulting in 270 control points and (d) a kernel width of 10.0 resulting in 1782 control points.

**Extended Data Fig 2.**
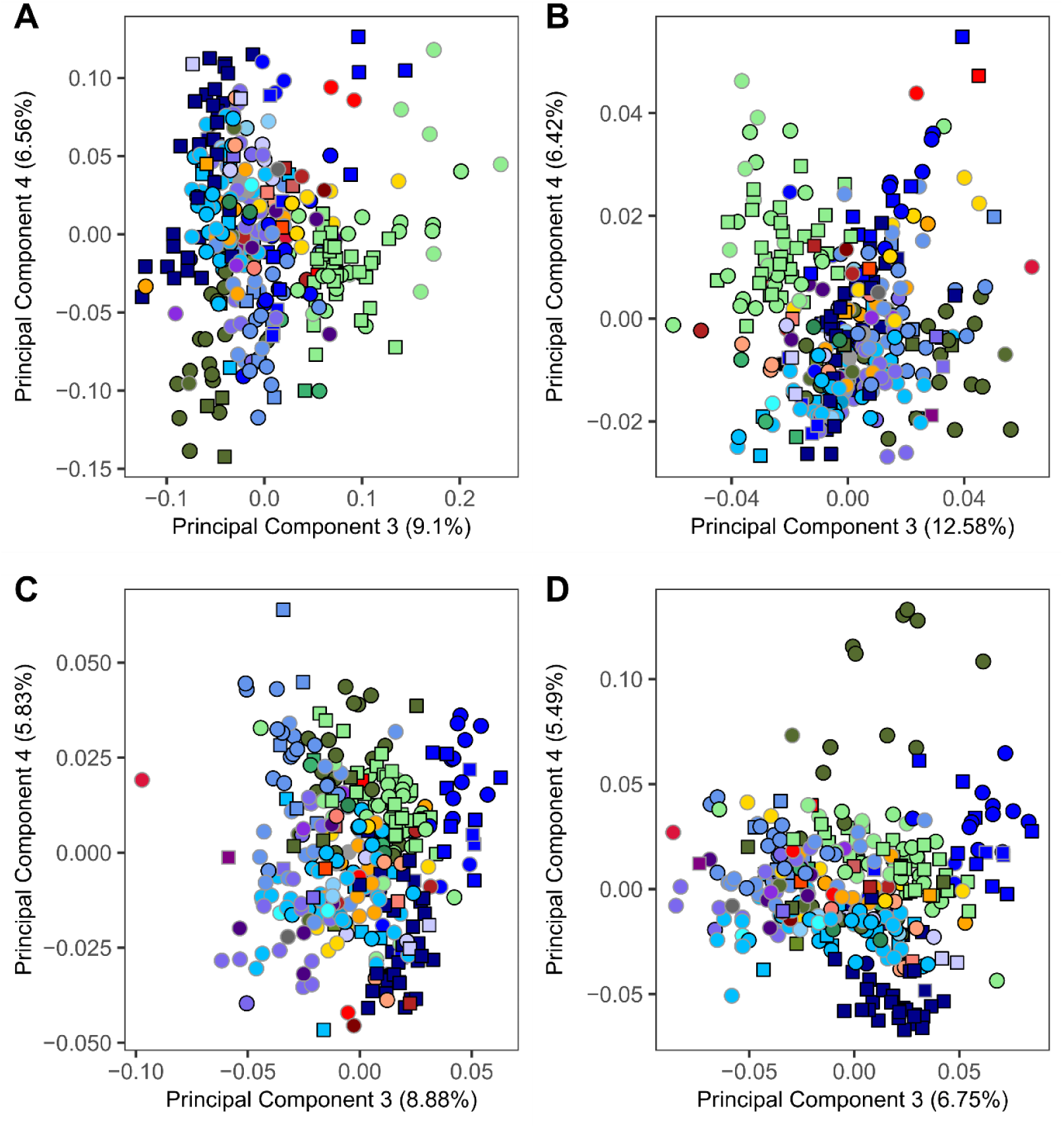
Comparison of manual landmarking and Deterministic Atlas Analysis (DAA) using Poisson meshes. Cranial morphospace for placental mammals with Poisson distributed specimens along axes PC1 and PC2, comparing the results between (a) manual landmarking with 754 landmarks and sliding semilandmarks and deterministic atlas analysis using (b) a kernel width of 40.0 resulting in 45 control points, (c) a kernel width of 20.0 resulting in 270 control points and (d) a kernel width of 10.0 resulting in 1782 control points.

## Extended Data Tables

**Extended Data Table 1.** Results of a two-way partial least squares (PLS) . Comparison of PC coordinates within morphospace obtained using manual landmarks to those obtained under the Deterministic Atlas Analysis (DAA). The results show there is a greater correlation as kernel width decreased (increased control points) and is significantly better with Poisson meshed compared to aligned-only specimens. All results are significant (p <0.001). The Effect Size (z) is listed in the brackets.

| Mesh Type | Measurement | 45 Control Points | 270 Control Points | 1782 Control Points |
| --- | --- | --- | --- | --- |
| Aligned-Only | r-PLS | 0.505 (4.86) | 0.537 (4.43) | 0.631 (2.90) |
| Poisson |  | 0.879 (8.74) | 0.887 (7.93) | 0.887 (5.07) |

**Extended Data Table 2.** Comparison of percentage variation controlled by each PC axes. Comparison of eigenvalues (percentage variation) obtained under manual landmarking and the Deterministic Atlas Analysis (DAA). The results show there is a greater correlation as kernel width increases (decreased control points) and is significantly better with Poisson meshed compared to aligned-only specimens. All results are significant (p <0.05).

| Mesh Type | Measurement | 45 Control Points | 270 Control Points | 1782 Control Points |
| --- | --- | --- | --- | --- |
| Aligned-Only | r | 0.9706 | 0.9807 | 0.9876 |
| Poisson |  | 0.9967 | 0.9887 | 0.9807 |

